# Diversity-disease relationships in natural microscopic nematode communities

**DOI:** 10.1101/2023.12.08.570776

**Authors:** Robbert van Himbeeck, Jessica N. Sowa, Hala Tamim El Jarkass, Wenjia Wu, Job Oude Vrielink, Joost A. G. Riksen, Aaron Reinke, Lisa van Sluijs

## Abstract

Biodiversity can affect parasite prevalence, a phenomenon widely studied in macroscopic organisms. However, data from microscopic communities is lacking, despite their essential role in ecosystem functioning and the unique experimental opportunities microscopic organisms offer. Here, we study diversity-disease effects in wild nematode communities by profiting from the molecular tools available in the well-studied model nematode *Caenorhabditis elegans*. Nanopore sequencing was used to characterize nematode community diversity and composition, whereas parasites were identified using nine distinct experimental assays based on fluorescent staining or fluorescent reporter strains. Our results indicate biotic stress is abundant in wild nematode communities. Moreover, in two assays, diversity-disease relations were observed: microsporidia and immune system activation were more often detected in relatively species-poor communities. Other assays, targeting different parasites, were without diversity-disease relations. Together, this study provides the first demonstration of diversity-disease effects in microbial communities and establishes the use of nematode communities as model systems for disease dilution.

## Introduction

Ecosystem functioning depends on biodiversity and biodiversity is shaped by parasites [1–3]. Increased (host) biodiversity can theoretically both cause increased (i.e. amplification effect) or decreased (i.e. dilution effect) parasite prevalence [2,4,5]. Generally, amplification effects occur when the parasite becomes more prevalent due to an increased availability of suitable hosts. Dilution effects results from reduced parasite reproduction because parasites experience difficulties finding a high-quality host amongst all species present. In contrast, parasites that replicate in a low-biodiversity population spread easily the because of high chances that the next organism they encounter is also a highly suitable host. Host-parasite relationships have been primarily studied in macroscopic species (e.g. [6–8]), but despite the necessity of well-functioning microscopic communities for ecosystem stability [3], natural host-parasite interactions in microscopic communities are little understood.

Interpretation of diversity-disease-correlations has not always been straightforward, as they depend on complex interactions within the natural community. Diversity-disease effects can be obscured or enhanced by parasite encounters, susceptibility and transmission of the species in the community, the order, timeframe and magnitude in which species enter or disappear from the community, life history traits of the host and host range of the parasite [4,7–21]. Interestingly, microscropic species that are relatively easy to grow and manipulate would present excellent models for studying diversity-disease effects, in particular when information from laboratory experiments can be combined with field observations. A model system where complex natural observations can be paired to controlled experiments will provide new experimental opportunities and biological insight into diversity-disease relations of (micro)organisms.

Nematodes present one of the most diverse and abundant groups of microscopic organisms worldwide [22] and some species possess characteristics that make them excellent model microscopic organisms. Of all nematodes, the free-living bacterivorous nematode *Caenorhabditis elegans* is a key model organism in many biological fields, ranging from developmental biology to neuroscience [23–26]. Today’s science not only focuses on the molecular and genetic understanding of this 1mm-sized organism in the laboratory, but considers its natural ecology too which led to the discovery of many naturally-infecting parasites [27]. Parasites infecting *C. elegans* and related bacterivorous species comprise opportunistic bacteria and obligatory parasites including oomycetes, microsporidia and viruses many of which cause deadly infections [27–32]. Recently, complete sequencing of the 18S gene using long-read technology of Nanopore was established as a technique to determine nematode community diversity and composition in large-scale experiments [33]. This enables high-resolution nematode community characterization, and thereby provides a new opportunity to study diversity-disease interaction in natural communities of the model nematode *C. elegans* [33].

In this study, we initially assess the (a)biotic environment where *C. elegans* was detected and thereafter, parasite presence was linked to the species diversity in nematode communities to investigate if diversity-disease correlates occur in microscopic organisms. During the autumn season natural nematode communities were collected from decaying organic matter. Sampling locations were chosen based on locations where *C. elegans* was previously found [34,35], so that molecular tools in this species could be used for efficient discovery of stress induced by bacterial, microsporidian and viral parasites. Morphological and Nanopore 18S gene amplicon sequencing reconstructed the nematode diversity and community networks which were connected to parasite observations.

Altogether, our results provide the first indication of disease-diversity relationships within microscopic communities and prospect a new experimental model system.

## Methods

### Sample collection and nematode extraction

Sampling was performed for five times during Autumn at the following three locations in the Netherlands: private garden (Heelsum), private vegetable garden (Wageningen), and patch of green surrounding a ditch (Renkum) (Supplementary Table S1). The active nematode community was extracted from the plant samples as previously described [36]. Additionally, details and a schematic overview covering the sample collection and nematode extraction are in Supplementary Text S1 and Supplementary Figure S1.

### Microsporidian identification in extracted nematode communities

Filtered nematodes were propagated for ~5 days on a 6cm NGM plate seeded with 10x *E. coli* OP50. Nematode communities that perished before this procedure or contained mites were not considered in subsequent analyses. Once the population was composed of a large quantity of non-starved adults, a portion of the plate was chunked onto a new seeded 6cm NGM plate to continually propagate potential infections. The remainder of the plate was then washed with 700 µl of M9 media and placed in a microcentrifuge tube. Samples were allowed to gravity settle for ~1-5 minutes, to remove contaminating bacteria and fungi through supernatant removal. 700 µl of acetone was added to the nematode pellet prior to spinning down in a centrifuge for 30 seconds at 8,000 rcf. The supernatant was discarded, and 500 µl of the chitin binding Direct Yellow 96 (DY96) solution (1x PBST, 0.1% SDS, 20 ug/ml DY96) was added. The samples were incubated for 30 minutes in the dark at 20°C prior to spinning down in a centrifuge for 30 seconds at 8,000 rcf. The supernatant was removed, and 20 µl of EverBrite Mounting Medium (Biotium) was added to the samples prior to mounting on glass slides for imaging using an Axio Imager.M2 (Zeiss). Z-stacks were captured at 63x using an apotome unit with maximum projection. Samples were considered infected with microsporidia if spore clusters were visible in a population. Spore sizes were assessed by measuring the length and width of at least 45 spores using Zen software.

To identify the species of microsporidia present within a nematode sample, molecular characterization was performed. Briefly, 20 large adult nematodes were placed in 10 µl of lysis buffer (50mM KCl, 100mM Tris-HCl, 2.5mM MgCl2, 0.45% NP40) and placed in a thermocycler at 65°C for 60 minutes, followed by 95°C for 15 minutes. 2 µl of lysate was then used as template in a PCR reaction with the forward primer V1F [5’ CACCAGGTTGATTCTGCCTGAC-3’] and the reverse primer 1492r [5’-GGTTACCTTGTTACGACTT-3’] or 18sR1492 [5’-GGAAACCTTGTTACGACTT-3’] to amplify microsporidian 18S. Sanger sequencing was performed on amplicons using the same primers listed above.

### Indication of present parasites or parasites in the homogenized substrate using fluorescent reporter strains

*C. elegans* reporter strains AGD926 (zcIs4[*hsp-4::GFP*]), SJ4100 (zcIs13[*hsp-6p::GFP*]) and AU133 (agIs17[*myo-2p::mCherry + irg-1p::GFP*]) were obtained from the Caenorhabditis Genetic Center (CGC). These strains indicate endoplasmic reticulum (ER) stress, mitochondrial stress or bacterial virulence respectively (Supplementary Table S2) [37] and were used to explore presence of harmful components in the microbiota of the homogenized substrate.

Starved populations from the reporter strains (AGD926, SJ4100, AU133) were transferred to fresh 6cm NGM plates containing *E. coli* OP50 just before nematode extraction from natural substrates. From each homogenized substrate 200µL blender solution was equally spread over the plates containing the reporter strains that were then incubated at 20ºC for 48h. After incubation, reporter nematodes exposed to blender solution were checked to investigate if GFP fluorescence was higher than GFP expressed by nematodes grown on control plates containing only *E. coli* OP50 [37]. Fluorescence was checked using an Olympus SZX10 microscope with a NIGHTSEA Microscope Fluorescence Adapter.

### Observation of intracellular parasites using Intracellular Pathogen Response and viral stress reporter strains

Extracted nematode communities were co-cultured with each of the following four reporter strains to observe activation of antiparasitic and antiviral pathways: SOW1 and SOW6 that detect upregulation of IPR genes *eol-1* and *pals-5* and SX2790 and SX2999 that detect upregulation of the antiviral gene *lys-3* (Supplementary Table S2) [38]. Fluorescence was checked once per day for 7 days using an Olympus SZX10 stereomicroscope equipped with an SZX-RFA stereo fluorescence illuminator unit and filter sets for GFP and RFP fluorescence viewing. Co-cultures were counted as positive if >10% of the reporter *C. elegans* were fluorescing.

### Extraction and screening for potential viruses

Co-cultures that resulted in reporter *C. elegans* fluorescence were washed off of test plates using M9 buffer and concentrated into 1 ml total volume via centrifugation. They were then homogenized via vortexing for 4min with 30-50 1 mm diameter silicon carbide beads (BioSpec Products). Homogenate was centrifuged to pellet debris, and supernatant was passed through a 0.22 µm pore size syringe filter (GenClone). Filtrates were seeded onto plates with SOW6, SOW1, SX2790, or SX2999 reporter strains and observed for fluorescence each day for 7 days.

### DNA and RNA extraction

DNA and RNA were isolated from samples that contained at least 100 nematodes at the moment of flash-freezing. From the 112 samples that met this requirement RNA and DNA was extracted as described previously by Harkes *et al*. [39]. In short, nematodes were bead homogenized using 3mm beads in a retch. Then RNA was isolated with a pH 4.5 phenol washing step, followed by a DNA isolation buffer washing step at pH 8.0. DNA and RNA concentrations were determined via Qubit fluorometer measurements (Thermo Scientific).

### Nematode DNA amplification

After DNA/RNA extractions, 81 samples contained sufficient DNA quantity for 18S rRNA amplification. Nematode communities were characterized using the workflow by van Himbeeck *et al*. (2024), with minor adjustments [33]. EXP-NBD196 (Oxford Nanopore Technologies ltd., UK) barcoded primers 988F (5’-ctcaaagattaagccatgc-3′) and 2646R (5′-gctaccttgttacgactttt-3′) were used to amplify the near-complete 18S rRNA fragment (~1750bp) [33,40]. PCR was performed in quadruplicate per sample and each reaction consisted of 12.5 µl LongAMP Taq 2x MM, 400 nM of each primer, 3 µl DNA template and 5.5 µl autoclaved Mili-Q water. After amplification all 4 PCR replicates were pooled and DNA amplification was verified on agarose gels. Finally, DNA concentrations were determined via Qubit 4 Fluorometer measurements (Thermo Scientific).

### Nematode 18S sequencing, read processing and species identification

The 18S rRNA was successfully amplified for 78 samples that were used for sequencing in four batches where each library contained equimolar ratios of all samples. Unwanted small fragments (<600bp) were removed prior to sequencing using NucleoMag NGS beads (0.5:1 bead:sample ratio) (Macherey Nagel). Library preparation was performed using the SQK-LSK112 kit, following the instructions of the manufacturer. Sequencing was performed on a MinION Mk1C using R9.4.1 flow cells.

Base-calling was performed using Guppy (v 6.2.1) in super accuracy mode (https://community.nanoporetech.com). The basecalled reads were demultiplexed using Guppy barcoder (v 6.2.1.) and adapters and barcodes were removed. Three samples (LvS26, LvS81, and LvS148) were excluded from the analysis at this stage because insufficient reads were obtained. Basecalling quality was controlled using NanoPlot (v1.40.0) (mean Phred score > 15) and Decona (v0.1.3) was used to filter 1400-2600bp reads with a >Q15 quality score [41,42]. Decona then clustered reads at 95% and Medaka consensus sequences were created from clusters with at least 100 reads. NCBI BLAST was used with an in-house database for taxonomic identification of nematode species [33,40]. Identifications with a similarity below 97% were excluded from the dataset. This data processing resulted in a OTU table containing the number of sequencing reads of each nematode taxon per sample. From this table, the number of species per sample was determined by summing the number of taxa with a read count larger than 0.

### Data analysis

All data was processed and visualised using the package *tidyverse* and *ggpubr* in custom written scripts in R (v 4.2.1) [43,44]. Generalized linear models (glm), Chi-square and Wilcoxon Rank Sum tests were computed with *R base*. The effect of abiotic factors on community compositions was tested after creating a ‘physeq’ object in the *phyloseq* package and performing a PERMANOVA based on Bray-Curtis dissimilarity index (n = 10,000) using ‘adonis2’ from the *vegan* package [45,46]. The *vegan* package was also used for calculating diversity indices (richness and Shannon index (H’)) with ‘specnumber’ and ‘diversity’ functions [45]. The nematode diversity was normalized (for substrate and community size) to facilitate cross-sample comparisons as the samples collected in this study varied because of ephemeral nature of the substrates and the consequent opportunistic sampling method that was applied. Therefore, normalized nematode richness was defined as the total number of nematode species (based on sequencing data) found in a sample, divided by the number of individuals (based on microscopy-based count data) per gram wet substrate found in that sample. Nematode co-occurrence networks were based on significant Spearman correlations (p adjust < 0.05) between species (that were collected at least three times) as calculated with the ‘rcorr’ function in the *Hmisc* package and adjusted by the Holm-Bonferroni method. Networks were visualized in Cytoscape (v3.9.1) when at least 3 species were present within a network.

### Data availability

*C. elegans* strains isolated in this study are available from CaeNDR (caendr.org) [34,35]. Custom written scripts and raw data files can be found on https://git.wur.nl/published_papers/van_himbeeck_2023_nematode_diversity_and_parasites/-/tree/main. The nematode sequencing data is accessible on SRA under BioProject PRJNA1021795. Processed data is included in the Supplementary Tables of this manuscript. Microsporidia 18S sequences are available in NCBI under the following accession numbers: OR636101, OR636102 and OR636103.

## Results

### Nematode community networks of the nematode C. elegans and co-occurring species

To link parasite presence to interspecies diversity in microorganisms, nematode communities were collected and analyzed. Nematode communities were obtained from decaying organic matter at three locations where *Caenorhabditis elegans* has previously been identified in the Netherlands [34,35] (Supplementary Figure S2, Supplementary Table S1). Nematode communities were subsequently extracted from the substrates, counted and the number of nematode species present was microscopically and molecularly established. Moreover, molecular analysis of the samples allowed for precise determination of the species present. The presence of *C. elegans* was of specific interest in this study, as this species was used as a reporter to monitor parasite presence.

In total 160 nematode communities were collected from ephemeral substrates and 76 of these were further characterised by sequencing (Supplementary Table S3). In these samples 58 different nematode species were molecularly characterised (Supplementary Table S4, S5). For 29 nematodes species only the genus name could be noted, as the used reference database does not always provide sequences annotated to species-level. The majority of the detected taxa were either bacterial or fungal feeders but also insect-parasitic, plant-parasitic and predatory nematodes were present (Supplementary Table S6). The estimated variance in nematode community composition originated for 20% from seasonal change, sampling location, moisture and pH differences (Supplementary table S7). Remaining variance in community compositions is likely in part attributable to biotic interactions, including nematode-bacteria, nematode-parasite, and nematode-nematode interactions [12,47]. Nematode-nematode interactions were examined by searching for specific cooccurrence or exclusion of certain species between samples. Most of the identified nematodes here were either opportunists or part of the basal fauna, although a few species exemplary for progressive succession (~5% of species) were also identified [48]. The species that formed cooccurrence networks were typically part of basal nematode fauna that is maintained over longer periods of time (Supplementary Table S4 and S8; Figure 1A). We did not observe any significant negative correlation between species. *Panagrolaimus subelongatus* (17), *Rhabditophanes sp*. (16), *Panagrolaimusrigidus* (16), *Acrostichus sp*. (15), *Rhabditoides sp*. (13), *Panagrolaimus sp*. (13), *Rhabditella axei* (12), *Pristionchus entomophagus* (10) and *Eucephalobus striatus* (10) co-occurred at least 10 times with *C. elegans* agreeing with previously found coincidences of these species (Félix and Duveau, 2012). Nevertheless, *C. elegans* nematodes did not coincide with these particular species more often than expected based on chance which underlines the opportunistic behaviour of *C. elegans* (Supplementary Table S4). Moreover, *Caenorhabditis* species are known to coexist together in some places [49–51]. Here, only one sample with a second *Caenorhabditis* species was obtained. This decaying kohlrabi contained *C. briggsae* and 7 other nematode species, but not *C. elegans*, despite molecular identification of *C. elegans* on other kohlrabies from the same location. *C. elegans* was detected in more than 71 % of the samples and we found that communities where *C. elegans* is present and absent have similar nematode richness (p=0.56, Wilcoxon Rank-Sum test). These findings support our use of it as a parasite reporter species for communities that differ in nematode diversity.

**Figure 1.**
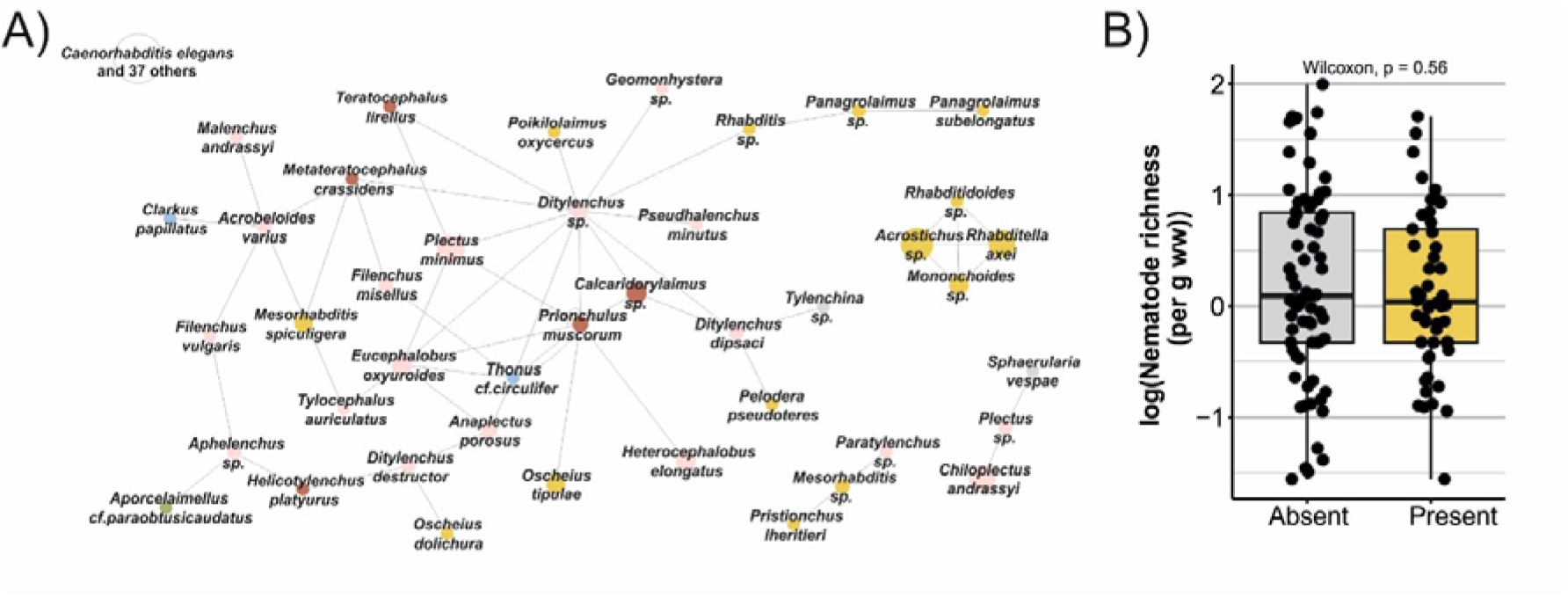
Nematode-nematode co-occurrences in decaying organic plants. A) Spearman-based (adjusted p < 0.05) co-occurrence networks of species that were present in at least 3 samples and contained at least 3 species. Colours indicate the c-p value of the species depicted (colonizer-persister value; higher values indicate species typically present in more mature nematode communities): yellow indicates c-p value 1, pink indicates c-p value 2, red indicates c-p value 3, blue indicates c-p value 4, green indicates c-p value 5, grey indicates c-p values are not known or variable for this genus [48]. Circle size reflects the number of samples in which a species was observed in the sequencing data. B) Nematode richness (in normalized species richness) in samples where *C. elegans* was absent or present.

### Nematode diversity determines parasite presence in natural field populations of nematodes

Collected nematode communities were screened for phenotypes that indicate: i) presence of microsporidia ii) mitochondrial stress, ER stress and stress caused by bacterial virulence [37], iii) IPR activation and iv) presumptive viruses [52] (Figure 2A). Some nematode communities died before they could be screened or could not be maintained successfully in the laboratory, and were excluded from further analysis (Figure 2B). All experiments harboured indications of parasite presence and 80% of the collected samples responded in at least one of the experimental assays (Figure 2C). In most cases, samples were found positive for multiple stressors (Figure 2C). Notably, these numbers likely underrepresent the total number of parasites occurring in the wild as not all may infect the reporter strains used, parasites might be lost during the extraction procedure and nematodes potentially evade parasites in some of the assays [53–55]. Taken together, these data suggest that biotic stress is a common burden for wild nematodes.

**Figure 2.**
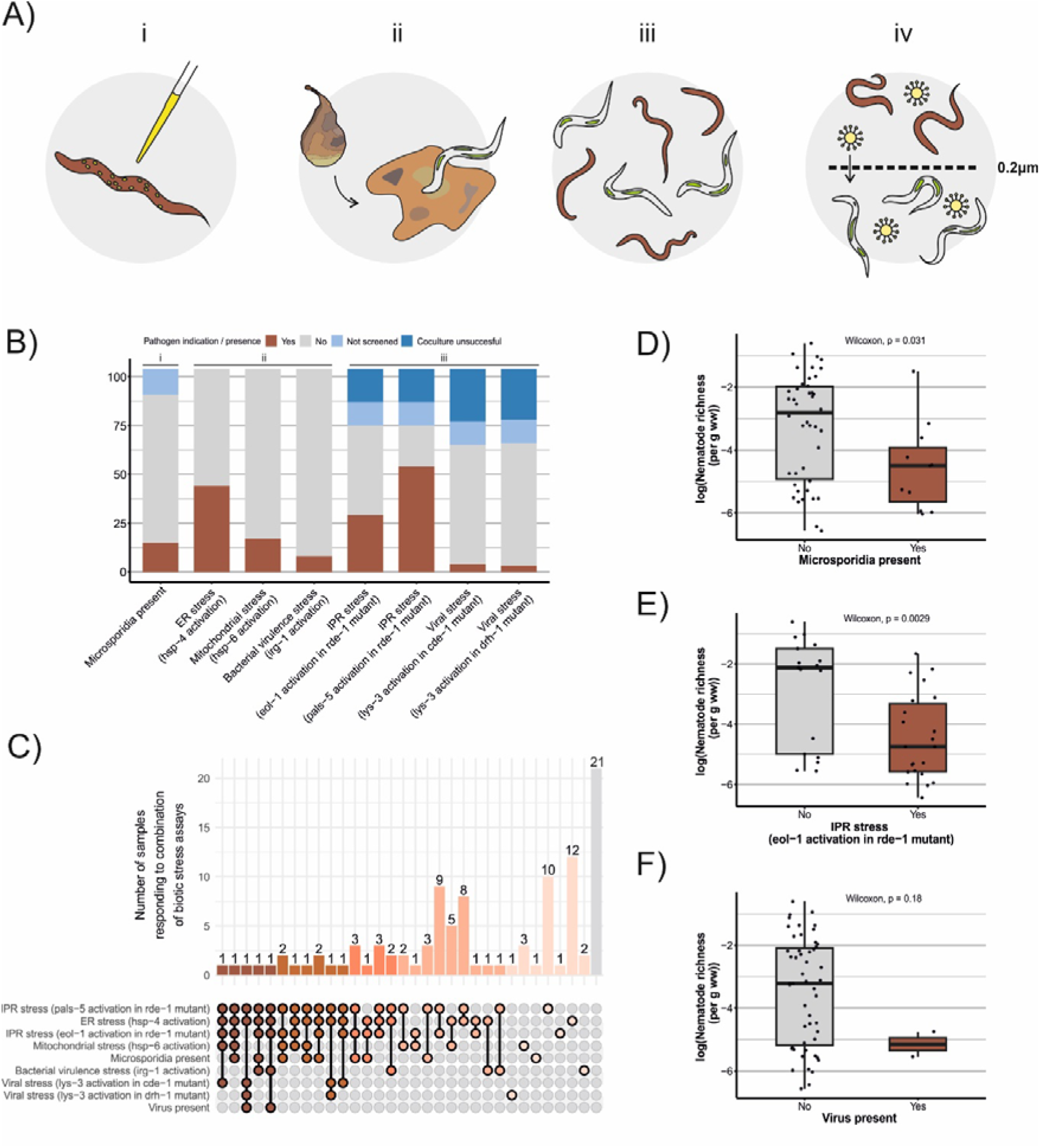
Biotic stress in nematode communities using *C. elegans* as a reporter species. A) The four different experimental assays to screen for biotic stress: i) staining of extracted nematodes to visualise microsporidia, ii) screening of *C. elegans* reporter strains on homogenized sample substrate containing a mixture of microbes, iii) co-culturing *C. elegans* reporter strains with the extracted nematode community and iv) co-culturing *C. elegans* reporter strains on 0.2 µm filtrate of the extracted nematode community B) The fraction of samples that was found to respond to each of the biotic stress indication assays. Because only a selection of samples was screened for virus presence after a 0.2 µm filter step (assay iv) these samples are not depicted here. C) Overlap of biotic stress indications within each of the samples. The plot shows how many samples (count y-axis) responded to specific combinations of biotic stress assays (x-axis). Colours illustrate the number of stress assays a sample was found positive for ranging from five (red) to none (grey). D) Normalized nematode richness for samples in which microsporidia were present or absent. E) Normalized nematode richness for samples which did or did not indicated mitochondrial stress on upregulation of the *C. elegans* IPR gene *eol-1* upon coculture. F) Normalized nematode richness for samples which did or did not contain *C. elegans* viruses.

Subsequently, we compared parasite presence to nematode diversity for each of the experimental assays (Supplementary Table S9). For two assays, diversity-disease correlations were observed: 1) microsporidia were more often present in communities with lowered nematode richness (Figure 2D) and 2) activation of the *eol-1*-based IPR reporter strain was more often observed in co-cultures when communities with a lowered nematode richness (Figure 2E). In agreement with previous studies, we found that presumptive *C. elegans*-infecting viruses were rare and observed in only 2 (1.5%) of the screened communities [31]. Both communities contained lower than average nematode richness (Figure 2F), but the low incidence of viral infections obstructs drawing solid conclusions from this observation. For the other experimental assays we did not observe diversity-disease relationships (Supplementary Figure S3, S4, Supplementary Table S3).

Diversity-disease relationships can be driven by host density, frequency or a combination of both [4,7,13,14]. To explore which of these may determine diversity-disease relationships in microbial nematode communities, the data gained in the microsporidian assay was explored further. We identified microsporidia in 15 out of 76 screened samples (16%) (Supplementary figure S5, S6). Almost half of these microsporidia isolates were found in *C. elegans* (Supplementary Table S10). The microsporidian isolates included the previously described species *Nematocida parisii* [28,56], but not all microsporidia could be molecularly determined and may represent novel species. Moreover, microsporidia were identified in *Pristionchus uniformus, Panagrolaimus rigidus* and in other nematodes of which the species could not be determined. Here, host density was defined as the number of host species per wet gram of substrate and host frequency was defined as the proportion of the host amongst the different nematode species in the sample. Analysis by a generalized linear model suggested a trend that host density (p = 0.058) as well as the interaction between host density and frequency (p = 0.072) might affect microsporidian presence in *C. elegans* (Supplementary Table S11), but additional data would be essential to observe if the same trend remains. As indicated above, key species may affect spread of parasites, but we could not pinpoint presence of key species in our relatively small dataset containing only 3 identified microsporidian hosts species (chi-square p > 0.05 for expected versus observed microsporidia infections per species). Microsporidia spores are known to be resistant to environmental conditions, including pH and desiccation [57,58]. Furthermore, pH has been shown to induce germination is thought influence the infectivity of microsporidia [59]. Here, microsporidia were more often observed on alkaline substrates, whereas substrate moisture varied (Supplementary Figure S7). In conclusion, because pH and nematode diversity were not correlated (glm, p = 0.46) (Supplementary Table S12), microsporidian presence in our samples was more likely to be observed in nematode communities with lower richness (possibly dependent on relatively higher host density- and frequency within the community) and an alkaline environment.

## Discussion

Although microscopic communities are vital for ecosystem functioning and stability, host-parasite relationships and the diversity-disease relationships have been poorly studied in microscopic communities. We argue that understanding diversity-disease effects in microbial communities is essential for predicting ecosystem functioning in a rapidly changing environment. Here we used wild nematode communities to study the effect of host diversity on the occurrence of parasites in microbial communities. Nanopore sequencing was applied for high-resolution characterization of nematodes present within each community. With this method we molecularly defined nematode communities naturally occurring with the important model organism *C. elegans*. Fluorescent reporter tools available in this model organism were used to identify bacterial-, microsporidian- and viral-induced stress inflicted on *C. elegans*. These screenings also discovered potentially new parasite species. In line with the dilution theory, some specialistic parasites were more often found in communities with fewer nematode species, indicating that diversity-disease effects can be found among microscopic organisms.

### Specific nematode-parasite interactions may underlie diversity-disease effects

Diversity-disease relationships within the nematode communities were tested via four different biotic stress assays examining nine classes of biotic stress. The vast majority of nematode communities in the collected samples responded in at least one of these tests, suggesting that biotic stress is frequent in wild nematode populations. Within these communities, we found correlation between nematode diversity and intracellular parasite prevalence as 1) samples containing microsporidian parasites had a lower nematode diversity and 2) the *eol-1-*based Intracellular Pathogen Response reporter strain more often responded to communities with a lower diversity. The experimental assays to indicate biotic stress covered diverse possible origins of stress-inducing parasites with varying host specificities. Parasites most prone to experience dilution effects are those that have a narrow host range [13]. Our data corresponds with this theory, as we observed potential dilution effects in microsporidian infected communities. Microsporidia are parasites of most types of animals with the vast majority of species only observed to display host and tissue specificity in one or two closely related hosts [59–61]. Microsporidia that infect nematodes also often display specialized tissue specificity, though there are examples of more generalist species [28,62]. Microsporidian occurrence in communities with lower species richness thus enhances their chance of encountering a high-quality host. Nevertheless, the lethal nature of microsporidia may also explain lower nematode diversity in communities suffering from microsporidia with a broad host range. We also observed potential dilution effects using the *eol-1* reporter strain in *rde-1* immunocompromised *C. elegans. Eol-1* is an RNA decapping enzyme that is upregulated during viral and microsporidian infections and is involved in activating enhanced RNAi in response to mitochondrial stress [63–66].

For other biotic stress reporter assays, we did not observed diversity-disease effects. The *hsp-6* reporter strain reports mitochondrial stress just like *eol-1*, but this reporter strain lacked the sensitivity-increasing *rde-1* mutation which is a possible explanation for not different observations between these two reporter strains. Also reporter strains (based on *lys-3*) that respond to viruses did not show a correlation with nematode diversity, despite the narrow host range of viruses [67]. As we have only identified two *C. elegans* viruses in our screening a larger study would be necessary to draw conclusions [31]. Another of the reporter strains responding to intracellular parasites (based on *pals-5*) did not show any diversity-disease relationship. It was noted that there were relatively few sequenced communities for which expression of this reporter strain was not observed. Finally, bacteria are usually opportunistic parasites not bound to certain hosts, therefore dilution effects were not expected, but amplification effects may occur. Nevertheless, these were not observed in the reporters strain that indicate bacterial virulence and also not in the ER stress reporter strain that detects stress caused by both generalists and specialists.

### Potential mechanistic bases of diversity-disease effects in ephemeral nematode communities

The short-lived ephemeral nematode communities sampled in this study consisted mostly of bacterial-feeding *r*-strategists that could experience strong competition for food in their temporary environment. The life history traits of the species present in these nematode communities may determine disease-diversity effects. Competition between species can drive dilution effects, especially in absence of predation [4]. Although several predatory species were observed, these were often present at low density and frequency. An exception was formed for facultative predators like *Pristionchus* [68], but these may also feed on bacteria in this habitat. Together, low predation and high competition circumstances suggest possibilities for observation of dilution effects, and we indeed observed a lower species diversity for nematode communities that were infected with microsporidia. Yet of note, potential predation by other species, such as mites or fungi was not studied here. Moreover, migration of ephemeral-specialised nematodes provides a spatial factor in the host-parasite dynamics that might be compared to that of a fragmented landscape. Fragmentation of ephemeral nematode communities may further contribute to the observation of diversity-disease effects [15,20,70,71].

### Detailed insight into ephemeral nematode community compositions that include C. elegans

To date, a comprehensive overview of nematode species co-occurring with *C. elegans* in its natural habitats was lacking. Historically, most studies investigated nematode community compositions in soil and water instead (e.g. for agricultural purposes or environmental monitoring). Here long-read sequencing data was used to identify community members that occur in the same habitats as *C. elegans*. We performed (nearly) full 18S sequencing to characterize the total nematode community on a low taxonomic level (mostly species or genus) [33], which yielded a total of 96 taxa. Subsequent studies may further broaden knowledge about *C. elegans* community compositions by simultaneously including other microfauna such as protists and bacteria. In agreement with previous studies, we identified nematodes that are known to co-occur with *C. elegans* from the genera *Caenorhabditis (C. briggsae), Oscheius (O. tipulae* and *O. dolichura), Panagrolaimus (P. detritophagus, P. rigidus, P. subelongatus), Rhabditophanes* and *Mesorhabditis (M. spiculigera)* and *Aphelenchoides* in our communities [49,50]. *E. striatus* is part of the basal nematode fauna, but all of the other species are species that quickly respond to enriched food resources [48]. The observation that these other nematodes do not specifically co-occur suggest that *C. elegans* can be considered an opportunist whose presence may be observed across suitable ecological niches of ephemeral substrates in the Netherlands.

### Conclusions

Biodiversity is important as it not only contributes to ecosystem functioning, but also to ecosystem stability, as exemplified by the diversity-disease relationships. Below-ground microbial communities - such as nematode communities - are vulnerable to shifts and biodiversity loss due to global change [72]. As microbial communities are vital for the majority of ecosystems, it is important to understand their response to a decrease in biodiversity in terms of parasite dynamics. Our study provides the first demonstration of diversity-disease relationships in microscopic nematode communities. Diversity-disease relationships depend on dynamic multifactorial processes in communities, and as such may be strikingly difficult to understand or capture. Studying diversity-disease effects in nematode populations poses a promising outlook for the future. In this study, we have extracted communities from the field through the use of a small plastic bag and have shown these could be observed and subsequently disassembled in the lab. Many collected species were successfully cultured afterwards, indicating follow-up experiments can be used to determine exactly which factors could cause diverse-disease effects to occur in nematode communities. Under laboratory settings, one can deduct or control factors such as species presence, density and select community members (including competitors and predators) based on specific life history traits. Furthermore, nematode parasites can be examined to trace infection and transmission (even in real time) [73,74]. Together, our results invite further characterization of host-parasite interactions in microscopic communities, and open opportunities for using nematodes communities as a diversity-disease model system in the lab and field.

## Acknowledgements

The authors thank Semih K. Aslan (primer design), Hans Helder (primer design, feedback on initial results), Stefan Geisen, Mark Sterken and Jan Kammenga (feedback on experimental set-up), Boas Kanis (sampling pilot), David Jordan and Eric Miska (reporter strains), Erik Andersen and Robyn Tanny (CaeNDR database) for their contributions. Some strains were provided by the CGC, which is funded by NIH Office of Research Infrastructure Programs (P40 OD010440). Funding for this research was provided by the Ecology Fund Grant of the Royal Netherlands Academy of Arts and Sciences. This work was supported by a Canadian Institutes of Health Research grant no. 400784 (to A.R.) and H.T.E.J was supported by a University of Toronto Open Fellowship and an Ontario Graduate Scholarship.

## Author contribution statement

LvS designed the research. RvH, JNS, HTEJ, WW, JOV, JAG, AR and LvS collected and processed nematode community, parasite and sequencing data. RvH and LvS performed computational analyses. RvH and LvS wrote the first draft of the manuscript, and all authors contributed to the interpretation of results and article writing.

## Supplementary information

**Supplementary Text S1 Sample collection and nematode extraction –** Detailed description of sample collection and nematode extractions.

**Supplementary Table S1 Summary of sampling location information** – The table indicates location details and number of collected samples at each location.

**Supplementary Table S2 *C. elegans* reporter strain genetic information**

**Supplementary Table S3 Collected sample data** – Contains information per sample collected including sample location, data, type, substrate weight, moisture, pH, nematode density and normalized species richness (based on morphological species identification) and whether or not *C. elegans* was isolated from the sample.

**Supplementary Table S4 Nanopore sequencing data output –** The data shows reads per species per sample.

**Supplementary Table S5 Sequencing abbreviations of nematode species –** The data shows reads abbreviations used for sequenced nematode species.

**Supplementary Table S6 Overview of molecularly identified nematodes in the sequenced nematode communities** The table indicates the nematode taxon, Maturity Index (according to its c-p value), feeding preference and percentage of the samples in which this taxon was discovered.

**Supplementary Table S7 Abiotic effects on community composition based on a permutation model**

**Supplementary Table S8 Co-occurring species in nematode communities** – Based on Spearman correlations and Hochberg-corrected p-values.

**Supplementary Table S9 Diversity indices of sequenced samples** – Shannon’s H, absolute richness (number of species) and normalised species richness (species corrected for sampling effort) for each sample based on sequencing.

**Supplementary Table S10 Data microsporidian species and hosts** – PCR-based host and microsporidian species identification and morphological characteristics of the nematode species identified. Some PCRs failed or nematodes were lost during the screening, here the missing data is indicated with NA.

**Supplementary Table S11 Effects of frequency and density on microsporidia of *C. elegans*** *–* GLM testing dilution and frequency effects on microsporidian occurrence in C. elegans.

**Supplementary Table S12 Correlation of normalised richness and pH –** GLM testing effect of pH on sequence-based normalised species richness.

