## Supplementary Text S1 for "Diversity-disease relationships in natural microscopic nematode communities"

**Supplementary methods**

*Sample collection and nematode extraction*

Three Dutch sampling locations were chosen, because these stably contained *Caenorhabditis* species for the last two to seven years in autumn(Cook *et al.* 2017). The locations were sampled five times every three weeks starting on August 30th 2021. At least 10 samples of decaying organic matter that appeared suitable habitats for bacterivorous nematodes (typically plants in a late stage of decay) were collected at each location. A similar sampling procedure was used for all trips (opportunistic selection of available decaying organic matter), but when possible previous finding locations of *C. elegans* or parasites of bacterivorous nematodes were resampled. Because of the perishable nature of the samples, resampling was not always possible. Additionally, on-site humidity and temperature were recorded, and samples were documented in five categories: fruits, vegetables, nuts, soil and (other) plant materials.

The Heelsum private garden has a partly forest-like nature with oak trees (*Quercus robur*) and undergrowth of dead-nettles (*Lamium* spp.) and green alkanet (*Pentaglottis sempervirens*) plants. Another part of the garden consists of a grass lawn with elevated vegetable garden boxes and a compost heap. The Wageningen vegetable garden contained about 15 different vegetable species and two compost heaps that are located adjacent to the vegetable patches. The edges of the garden contain several trees and fruit scrubs. The Renkum patch of green mostly contains butterbur (*Perasites hybridus*) plants and common nettle (*Urtica dioica*) surrounding a ditch.

The active nematode community was extracted from the plant samples as previously described (Van Bezooijen 2006). In short, each sample (1-20g) was homogenized in a ~100mL of water using a kitchen blender (Princess, model 2005) at full speed for 5-10s. The pH of 2mL homogenized blender solution was measured at room temperature (Jenway pH electrode 924 904). Moreover, 200µL was placed on each of the three 6cm NGM plates containing either the *C. elegans* reporter strain AGD926, SJ4100 or AU133 (see bacterial reporter strain assay below). The remaining blender solution was placed on a custom-made nematode filter consisting of a Universal Hygia Milk filter (928157-56) placed on top of a Lekko (S475-30) filter clamped in a nematode extraction sieve wherein nematodes were extracted overnight in ~100mL water at room temperature (~20ºC). For each sample, dry weight and substrate moisture were determined after drying a subsample (1-10g) for 48h at 60ºC.

After nematode extraction, the solution containing the nematodes was put in a glass beaker and left to settle for 1h. Next, two-thirds of the supernatant was discarded and the remaining solution was transferred to a glass centrifuge tube. After the solution settled for 1h, a 1mL pellet (containing the nematodes) was transferred to a 9cm nematode growth medium (NGM) plate without bacteria. Then the number of nematode species in the population was determined morphologically under a Leitz Wetzlar stereo microscope (based on adult female characteristics including size, vulval placement, mouth and tail structure) and any remarking population characteristics were noted (e.g. presence of unhealthy looking nematodes). A maximum of three potential *C. elegans* nematodes per plate were isolated for further propagation and species-specific PCR test (Barrière & Félix 2005). Half of the isolated wild nematode community was flash-frozen in liquid nitrogen 48h after the start of the nematode extraction for sequencing purposes. The remaining 50% of this isolated community of nematodes was split in two samples of 25% and placed on 9cm plates containing *E. coli* OP50. These samples were subsequently used for microsporidian and virus identification (see Methods). Some populations containing few nematodes were not split and the full sample was flash frozen for sequencing instead. These samples were not considered in further analyses of parasite presence.

Samples were kept at 4ºC from sampling onwards, except for the nematode extraction, species diversity check and selection of nematodes for propagation that were performed at room temperature (~20ºC). Nematode extraction took 22h on average and screening and selection of the nematodes added about another 4h at room temperature.
