## Supplementary Figure S1-S7 for "Diversity-disease relationships in natural microscopic nematode communities"

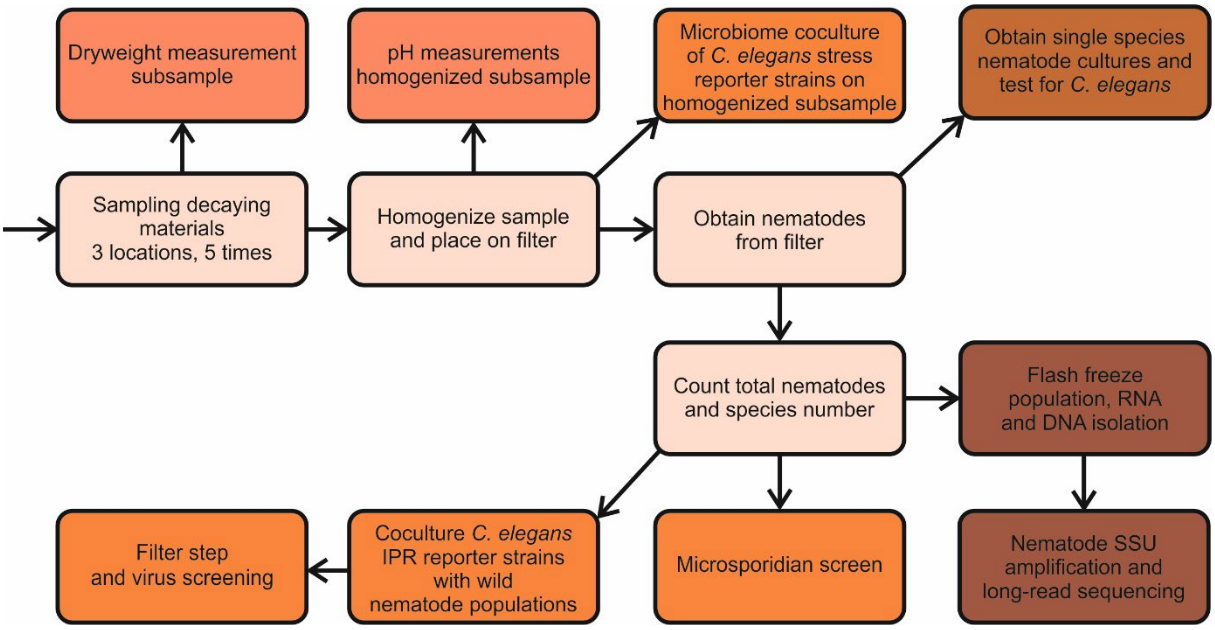


**Figure S1 Experimental workflow of the overall study** – Nematodes were collected from three different samples sites on 5 separate occasions in autumn. Of these the dry weight and pH were determined. After the nematode populations were collected, single nematodes were singled out and when applicable tested for presence of *C. elegans*. Three methods were applied to screen for abiotic stresses: co-culturing of reporter strains with the homogenized subsample, co-culturing of a subsample of the extracted nematodes with reporter strains, and direct staining of a subsample of the extracted nematodes to visualize microsporidia. Finally, the remaining nematodes were flash frozen to isolate RNA and DNA. The nematode SSU in these samples was PCR amplified and long-read sequencing was performed to identify nematode species present in the samples.

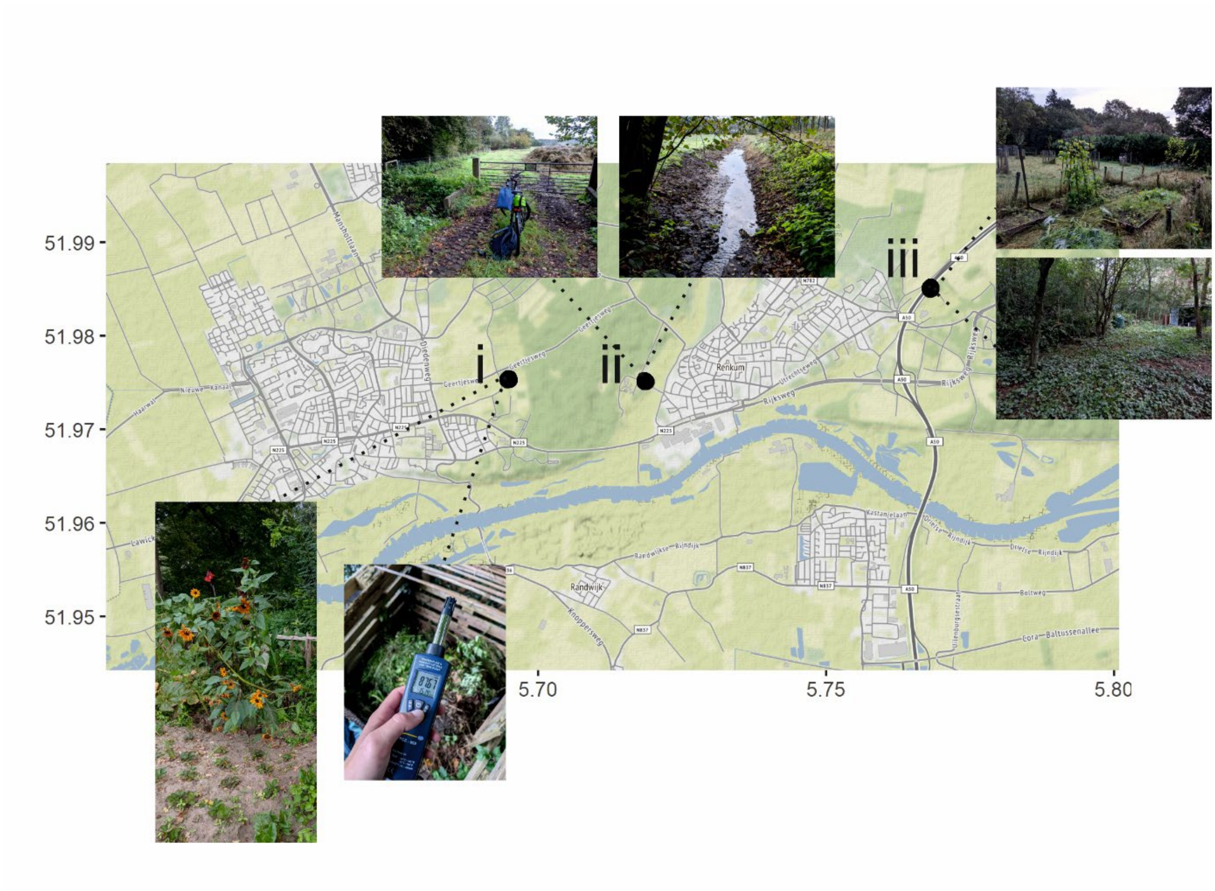


**Figure S2 Sampling locations –** Samples were collected around Wageningen at three locations: i) a vegetable garden, ii) a patch of green surrounding a ditch and iii) a private garden covering a vegetable garden and forest edge. Axis labels indicate GPS coordinates (x-axis: longitude; y-axis: latitude).

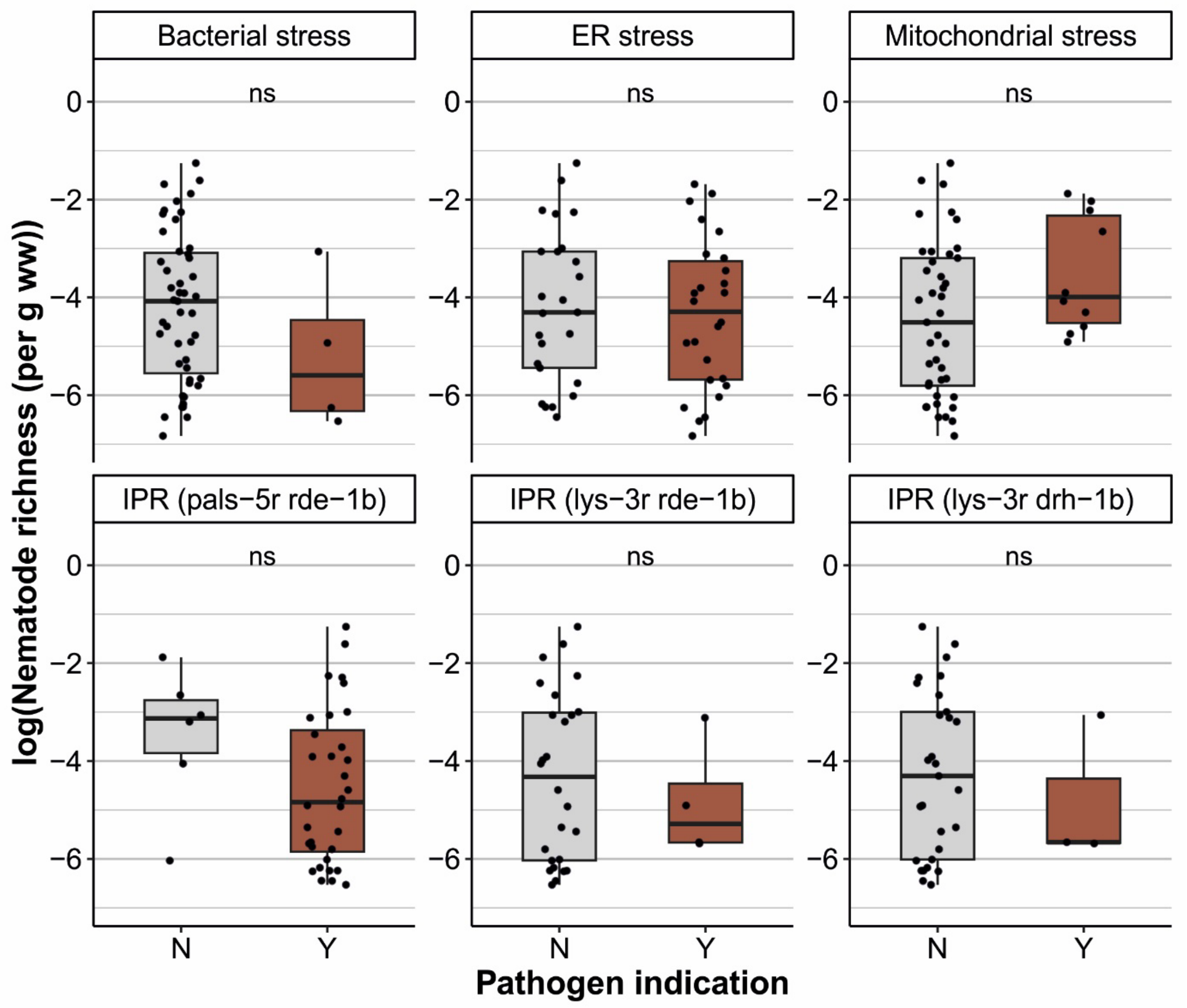


**Figure S3 Correlation plots of normalized nematode richness where parasite presence did not correlate significantly** – Sequence-based normalized nematode richness for samples which did or did not indicated bacterial stress (upregulation of the *C. elegans* gene *irg-1*), ER stress (upregulation of the *C. elegans* gene *hsp-4*), mitochondrial stress (upregulation of the *C. elegans* gene *hsp-6*) or Intracellular Pathogen Response (IPR) stress (by upregulation of the *C. elegans* genes *pals*-5 or *lys*-3).

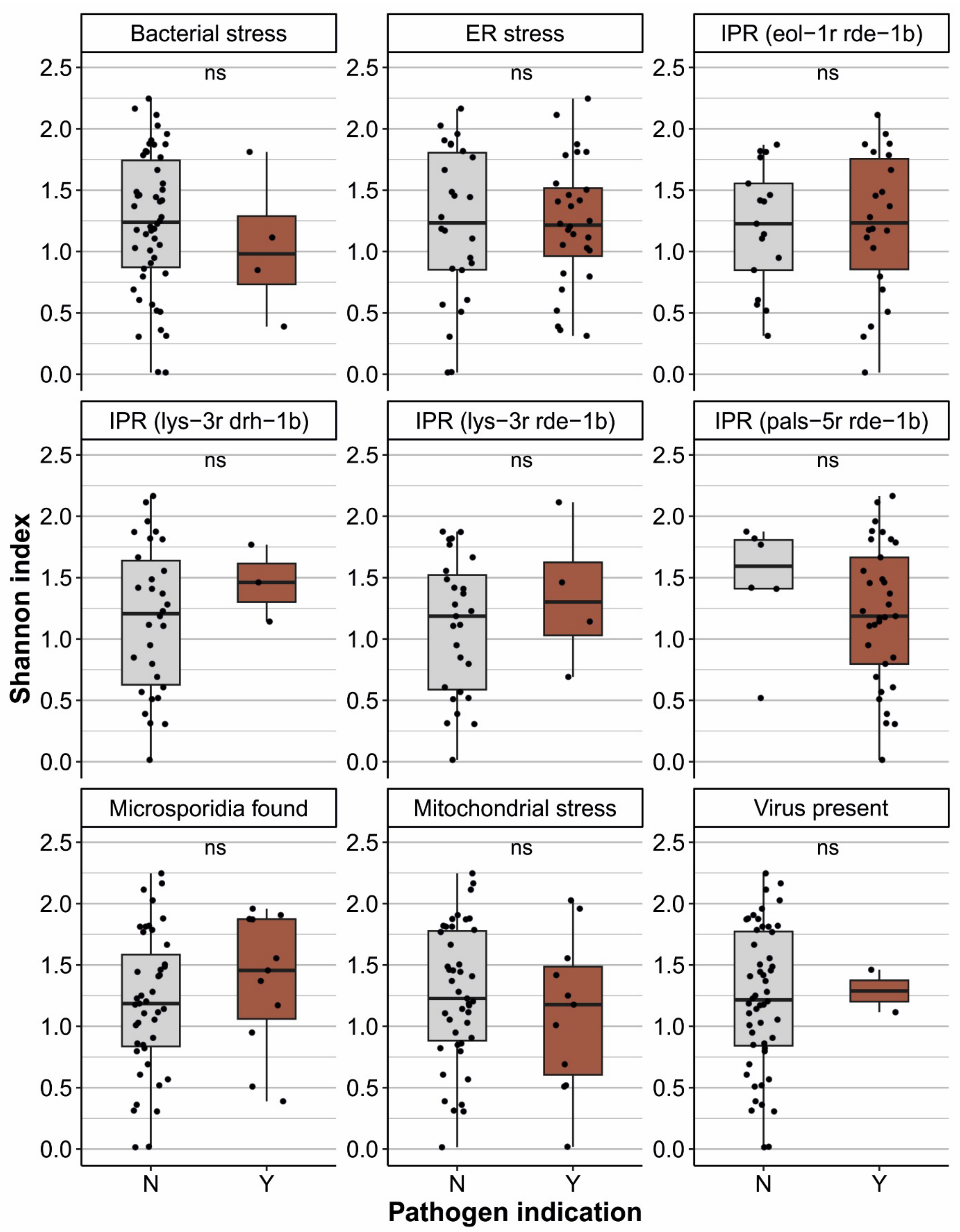


**Figure S4 Shannon diversity and parasite presence –** Sequence-based Shannon diversity for each of the nine performed screenings that indicate parasite presence in wild nematode populations. Plots indicate if samples did (Y) or did not (N) indicate bacterial stress (upregulation of the *C. elegans* gene *irg-1*), ER stress (upregulation of the *C. elegans* gene *hsp-4*), mitochondrial stress (upregulation of the *C. elegans* gene *hsp-6*) or Intracellular Pathogen Response (IPR) stress (by upregulation of the *C. elegans* genes *pals*-5, *eol-1* or *lys*-3) or presence of viruses or microsporidia.

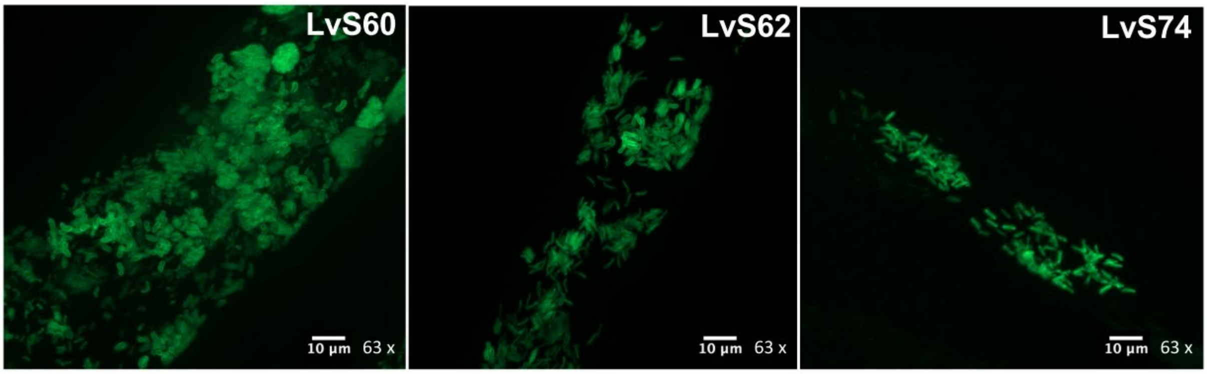


**Figure S5 Microsporidia infection** – Nematodes stained with DY96 and imaged at 63x. The 18S region of these samples were successfully amplified via PCR.

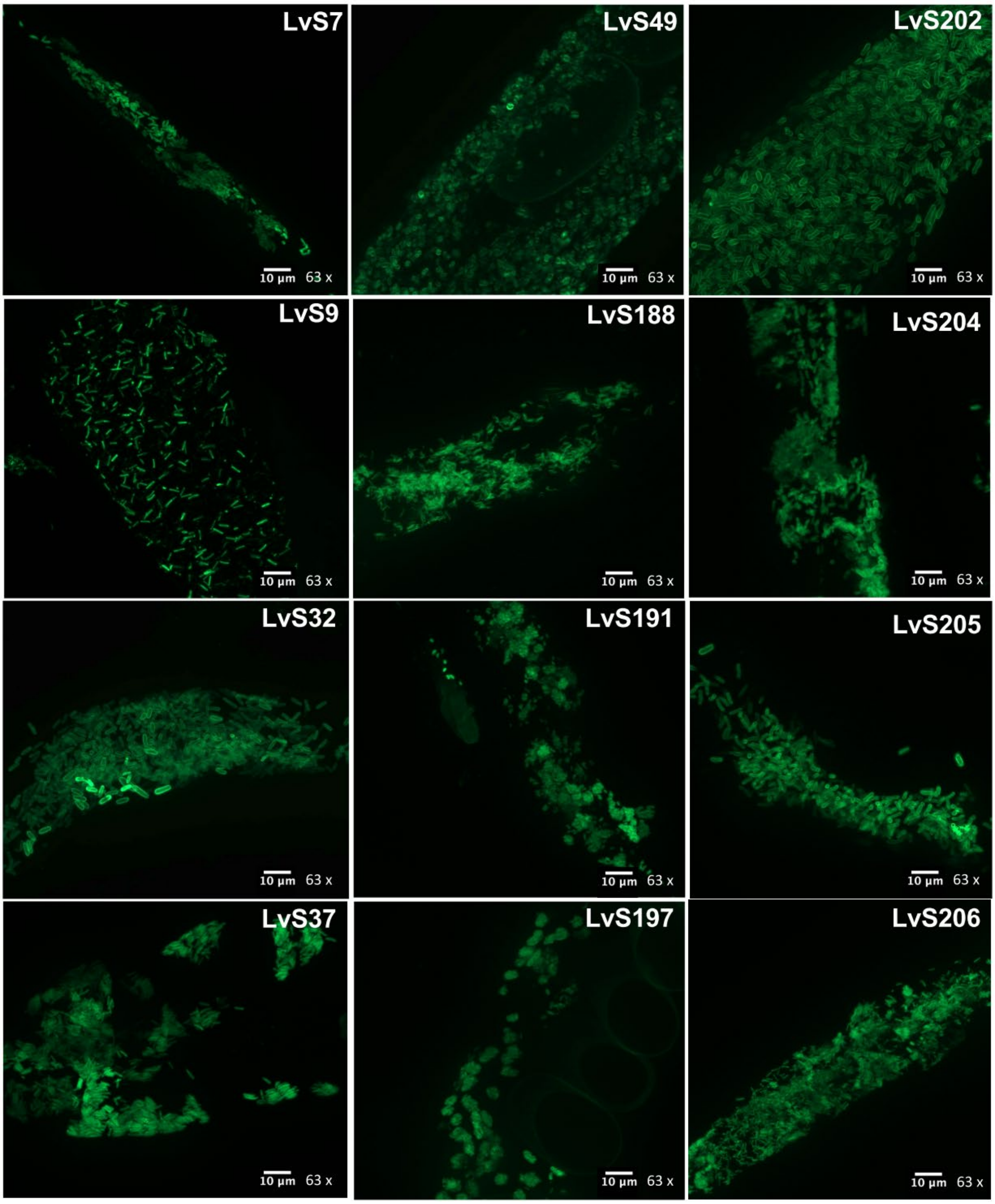


**Figure S6 Microsporidia infection** Nematodes stained with DY96 and imaged at 63x. The 18S region of these samples was not successfully amplified by PCR.

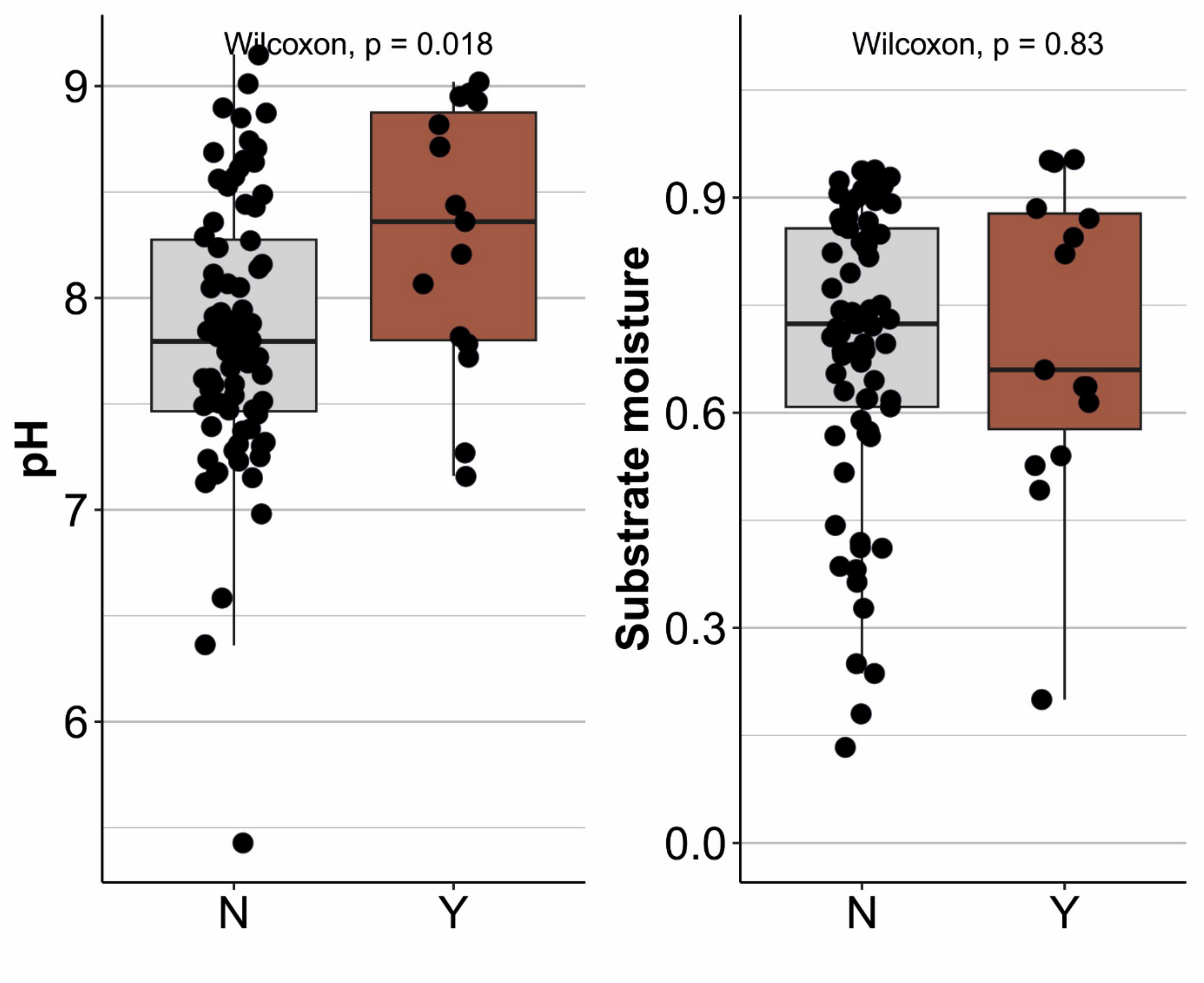


**Figure S7 Abiotic characteristics of samples screened for microsporidia –** Substrate pH and substrate moisture are shown for samples that did (Y) or did not (N) contain microsporidia.
